# PepSeA: Peptide Sequence Alignment and Visualization Tools to Enable Lead Optimization

**DOI:** 10.1101/2021.10.26.465927

**Authors:** Javier L. Baylon, Oleg Ursu, Anja Muzdalo, Anne Mai Wassermann, Gregory L Adams, Martin Spale, Petr Mejzlik, Anna Gromek, Viktor Pisarenko, Dzianis Hancharyk, Esteban Jenkins, David Bednar, Charlie Chang, Kamila Clarova, Meir Glick, Danny A. Bitton

**Author notes:** **Corresponding Authors, Javier L. Baylon**, Modeling and Informatics, Merck & Co., Inc. 33 Avenue Louis Pasteur, Boston, MA 02115, USA, **Danny A. Bitton**, Na Valentince 4, FIVE Building, Prague 5 - Smichov, Prague, 150 00, Czech Republic.

## Abstract

Therapeutic peptides offer potential advantages over small molecules in terms of selectivity, affinity, and their ability to target “undruggable” proteins that are associated with a wide range of pathologies. Despite their importance, there are currently no adequate molecular design capabilities that inform medicinal chemistry decisions on peptide programs. More specifically, SAR (Structure-Activity Relationship) analysis and visualization of linear, cyclic, and cross-linked peptides containing non-natural motifs, which are widely used in drug discovery. To bridge this gap, we developed PepSeA (**Pep**tide **Se**quence **A**lignment and Visualization), an open-source, freely available package of sequence-based tools (https://github.com/Merck/PepSeA). PepSeA enables multi-sequence alignment of non-natural amino acids and enhanced HELM (Hierarchical Editing Language for Macromolecules) visualization. Via stepwise SAR analysis of a ChEMBL peptide dataset, we demonstrate PepSeA’s power to accelerate decision making in lead optimization campaigns in pharmaceutical settings. PepSeA represents an initial attempt to expand cheminformatics capabilities for therapeutic peptides and to enable rapid and more efficient design–make–test cycles.

## INTRODUCTION

Naturally occurring peptides possess powerful therapeutic properties given their inherent ability to selectively bind to molecular targets and elicit a desired biological response. Peptides exert important physiological functions as hormones, growth signals, neurotransmitters, quorum sensing, and anti-infective agents, holding a great promise for the treatment of various human disorders including metabolic, neurodegenerative, cancer, and infectious diseases.^1^ Over the last few decades the advent of recombinant protein technology, along with the development of protein purification methods and solid-state peptide synthesis, has facilitated a more widespread use of protein and peptide therapeutics. Many recombinant human proteins and peptides have been approved as medicines. Examples include monoclonal antibodies for the treatment of cancer, autoimmune and infectious diseases,^2–5^ protein vaccines,^6,7^ synthetic peptides, and modified natural enzymes.^8–11^ To date, there are around 80 peptide drugs on the global market,^12^ and this figure is likely to grow given the increased investment and research in the peptide field. Antimicrobial peptides, acting against a wide range of pathogens, show great promise as next-generation antibiotics.^13^ Another promising area in medicinal research towards targeted therapeutics is the development of peptide-drug conjugates, comprised of a cytotoxic payload linked to a peptide moiety that targets a protein receptor overexpressed in cancer cells.^14^

Peptides are biopolymers composed of up to around 40 amino acids. Because of their size (molecular weight ranging between 500 – 5000 Da) they are placed somewhere between smallmolecule drugs and biologics. As such, peptides do not follow Lipinski’s rule of 5 (Ro5),^15^ traditionally considered to predict the “drug-likeness” of a molecule. Most of the orally available peptides in clinical use are up to 1200 Da in molecular weight (MW), have partition coefficient values in the −5 to 8 range, and have 5 times more hydrogen-bond donor and acceptor groups than is prescribed by the Ro5.^16^ In addition, clinical peptides are more flexible than traditional small molecule therapeutics in terms of the number of freely rotatable bonds and also contain total polar surface area (TPSA) well over the value required for passive membrane permeability (< 140 A^2^).^16^ These properties beyond Ro5 (bRo5) have led to new computational strategies for peptide design. For example the use of conformationally dependent descriptors, such as radius of gyration for size or 3D-polar surface area and desolvation free energies for polarity, can better predict the passive permeability of bRo5 molecules, including peptides.^17^ A key advantage of peptides over smallmolecule drugs is their increased structural complexity and balance of conformational flexibility and rigidity, which enables them to bind more selectively and with high affinity to their protein targets. Peptide ligands can also bind to extended and shallow protein surfaces, making them ideal for targeting the “undruggable” targets, such as protein-protein interactions (PPIs) which are associated with a wide range of pathologies.^18^

The goal of peptide drug design is to generate optimized peptide leads that couple target affinity with favorable target access properties (i.e., cell-entry ability for intracellular targets) and metabolic stability. In the absence of structural information of protein-peptide complexes to guide rational peptide design, the diverse sequence space of peptides and their non-natural analogs can be sampled by screening for affinity using natural or synthetic peptide libraries^19^ and techniques such as phage display^20,21^ and mRNA display.^22–24^ Membrane permeability, a key predictor of oral bioavailability, can be improved by introducing modifications to the amide backbone employing approaches such as N-methylation,^25,26^ incorporation of non-canonical amino acids and non-peptidic fragments,^27^ or macrocyclization.^28^ Proteolytic stability of peptides has been shown to increase with macrocyclization^28,29^ as well as by the introduction of N-terminal modification, such as acetylation and methylation. Additionally, incorporation of D-amino acids, thioamides, and other amide bond mimetics have been shown to improve peptide stability.^12^ Herein is explained how the data generated by these and other approaches is leveraged to build robust structure-activity relationships (SAR) in order to design new peptide hits during the design cycle.

Optimizing peptide hits into drugs remains a significant challenge for the medicinal and computational chemistry community. While the field of cheminformatics is well established for small molecules, there are no adequate solutions for peptide SAR analysis and for visualization of the linear, cyclic, and cross-linked peptides that contain non-natural motifs and are widely used in pharmaceutical research. For example, the depiction of bicyclic peptide structures is generally challenging and not intuitive for simple visual inspection. This manuscript presents PepSeA (**Pep**tide **Se**quence **A**lignment and Visualization), an open-source package of sequence-based tools that overcomes visualization challenges for peptide SAR and addresses this unmet need by employing the Hierarchical Editing Language for Macromolecules (HELM).^30^ More specifically, we describe how PepSeA tools can be used for:

1. Multi sequence alignment of non-natural amino acids to enable assay data comparison for multiple peptides.
2. Positional SAR analysis that leverages the sequence representation of peptides using HELM.
3. Deployment of API-based tools to enable SAR exploration and visualization.

The tools developed in this study are open source and freely available for use (https://github.com/Merck/PepSeA). These capabilities became an integral part of the peptide design cycle at our pharmaceutical company. Biopolymer registration using HELM notation ensures that both the peptides and their bioactivity data are model-ready and outlive the program. The use of HELM and PepSeA tools accelerate SAR elucidation in peptide projects and enable medicinal chemists to better understand the relationships between calculated molecular properties and various experimental endpoints of interest, including affinity, permeability, solubility, and stability of peptides. In this work, the functionalities of the different PepSeA components are described. Also presented is a data analysis workflow example, using a publicly available dataset from the ChEMBL database, that one can prospectively apply to drug discovery programs.^31,32^

## METHODS

### Preparation of ChEMBL Peptide Dataset

To showcase PepSeA, a dataset of peptides with HELM notation and activity data was extracted from the ChEMBL database^31,32^ as follows: First, compounds with four consecutive amino acids in their full atomistic structure were extracted from the database (34998 compounds). This step was performed to identify peptide-like compounds that might not be annotated with HELM and that could be converted to HELM using the new rules as described below. After this step, molecular data for each of the filtered compounds were acquired using the ChEMBL web services to obtain HELM representation of compounds and activity data if available. This process resulted in a set of 15259 peptides with HELM sequences. The HELM sequences and activity data were next aggregated, and the following filters were applied: organism = “*Homo sapiens*” and number of unique compounds with activity data for a given ChEMBL assay id >= 10. This filtering resulted in a dataset comprised of 429 subsets of peptides that had been tested against the same target in the same assay. Finally, a few of these subsets were selected to test the tools. For sequence alignment examples, those subsets that had the largest number of peptides were selected, i.e., assays CHEMBL1008221 (256 peptides), CHEMBL956457 (151 peptides), and CHEMBL660105 (150 peptides). For the data analysis workflow example, subset CHEMBL1819839, which corresponds to previously published activity data of vasopressin analogues, was selected.^33^

Additionally, HELM sequences for 300 peptidic molecules (with four consecutive amino acids) were generated using the new HELM rules to represent click and Ring-Closing Metathesis (RCM) chemistries as described in the Results section. The datasets are provided in the Supplementary Information.

### Multiple Alignment using Fast Fourier Transform (MAFFT) for HELM Peptide Sequences

To implement the sequence alignment described in this study, the latest version of the Multiple Alignment using Fast Fourier Transform (MAFFT) program was employed.^34,35^ The details of the method and its improvements over time have been extensively reviewed and described elsewhere.^34–39^ Briefly, MAFFT is a multiple sequence alignment program that uses the fast Fourier transform to rapidly identify homologous segments in sequences.^34^ For the implementation of sequence alignment with MAFFT, two key functionalities of the program were used: the ability to use extended alphabets (i.e., beyond 20 letters of natural amino acids) and support for custom substitution matrices for alignment. MAFFT allows alignment of sequences containing 248 different ASCII characters.^40^ For each alignment job, a lookup table was created from the HELM symbols of all amino acids occurring in the input sequences into a custom alphabet of 248 characters, therefore allowing at least 228 different non-natural amino acids in one alignment. The same mapping is also applied to a custom substitution matrix. To perform sequence alignment of peptides with non-natural amino acids, each HELM input sequence is preprocessed and converted to its ASCII representation using the lookup table. After the alignment is performed the sequences are converted back to the original HELM using the same lookup table, creating a file with an extended FASTA format that uses multi-character residues corresponding to amino acid HELM symbols.

### Substitution Matrix Calculated with Rapid Overlay of Chemical Structures (ROCS) Similarities Across ChEMBL 28 HELM Monomers

To support the alignment of peptides with non-natural amino acids using MAFFT, a custom substitution matrix was generated using HELM monomers in the ChEMBL 28 library. To generate the matrix, the subset of 779 monomers found in the HELM peptide dataset described in the previous section was used. The generated substitution matrix format is similar to BLOSUM matrix^41^ with the scores calculated based on a chemical similarity between amino acids. The steps to generate the matrix are as follows (Figure S1): (i) generate a preset number of conformers for each amino acid using Omega2;^42^ (ii) calculate ROCS^43^ score between each amino acid and conformer pair; (iii) select the best Tanimoto Combo score for each amino acid pairs and rescale the score to [-10,10] range. The substitution matrix is provided in the Supporting Information.

### HELM Visualization Tools

The visualization tools presented consist of two main components: 1) A visualization library and 2) A visualization API server. The components are written in JavaScript. A description of the components is provided in Supplementary Information. Briefly, the visualization library is a module for transforming a HELM string to an SVG object. Finally, the visualization API server is a web-service based on the visualization library that generates and returns HELM depiction.

## RESULTS AND DISCUSSION

### Cyclic Peptides: Extending HELM Beyond Disulfide and Amide bonds

An expansion of the HELM notation was performed to accommodate the complex connections often found in macrocyclic peptides, comprised of both natural and non-canonical amino acids, through an assignment of additional R groups to define attachment points on monomers. Frequently used synthetic linkages such as triazoles and alkenes, resulting from click^44^ or RCM^45^ reactions respectively, move beyond the disulfide and amide bonds found in nature and require special treatment. The new R groups that have been added are Vinyl (C=C), Azide (N+[N+]=[N-]), and Alkyne (C#C). These new capping groups correspond to the reactive moieties that are involved in the above-mentioned reactions.^44,45^

HELM is a notation that describes final products without any information concerning how they are made, or which reagents are used, but assigning these capping groups allows a chemist to use the same PEPTIDE sequence without having to change any monomers to describe the structures both before and after the crosslink is achieved synthetically. Figure 1a shows some simple examples of PEPTIDE monomers that use the new R groups. These new R groups are used on monomer sidechains and are linked using CHEM monomers that capture the resulting product motif when the linkage is formed. For products of RCM reactions, 3 new monomers were added (**Figure 1**b). Peptides that have trans double bonds use the tRCM monomer, and those with cis double bonds use the cRCM monomer. For the RCM monomers Cl was chosen for the R group caps, but any R groups could be used, since both R groups are always connected to another chain in the products. When the peptide being described is a mixture or the stereochemistry is unknown, the xRCM monomer can be used along with 3rd section annotation to indicate whether the peptide is a single isomer or a mixture. For products of click reactions, two new monomers, [14Triazole] and [15Triazole], have been added to describe the two possible regioisomers that can result from the cycloaddition (**Figure 1**c). Using these new caps and CHEM monomers, the generation of HELM sequences for an expanded set of peptides is possible (**Figure 1**d). The use of these new rules to convert more cyclic peptides to HELM notation can facilitate the sequence-based analysis of compounds and their activity with the tools described in the next sections.

**Figure 1.**
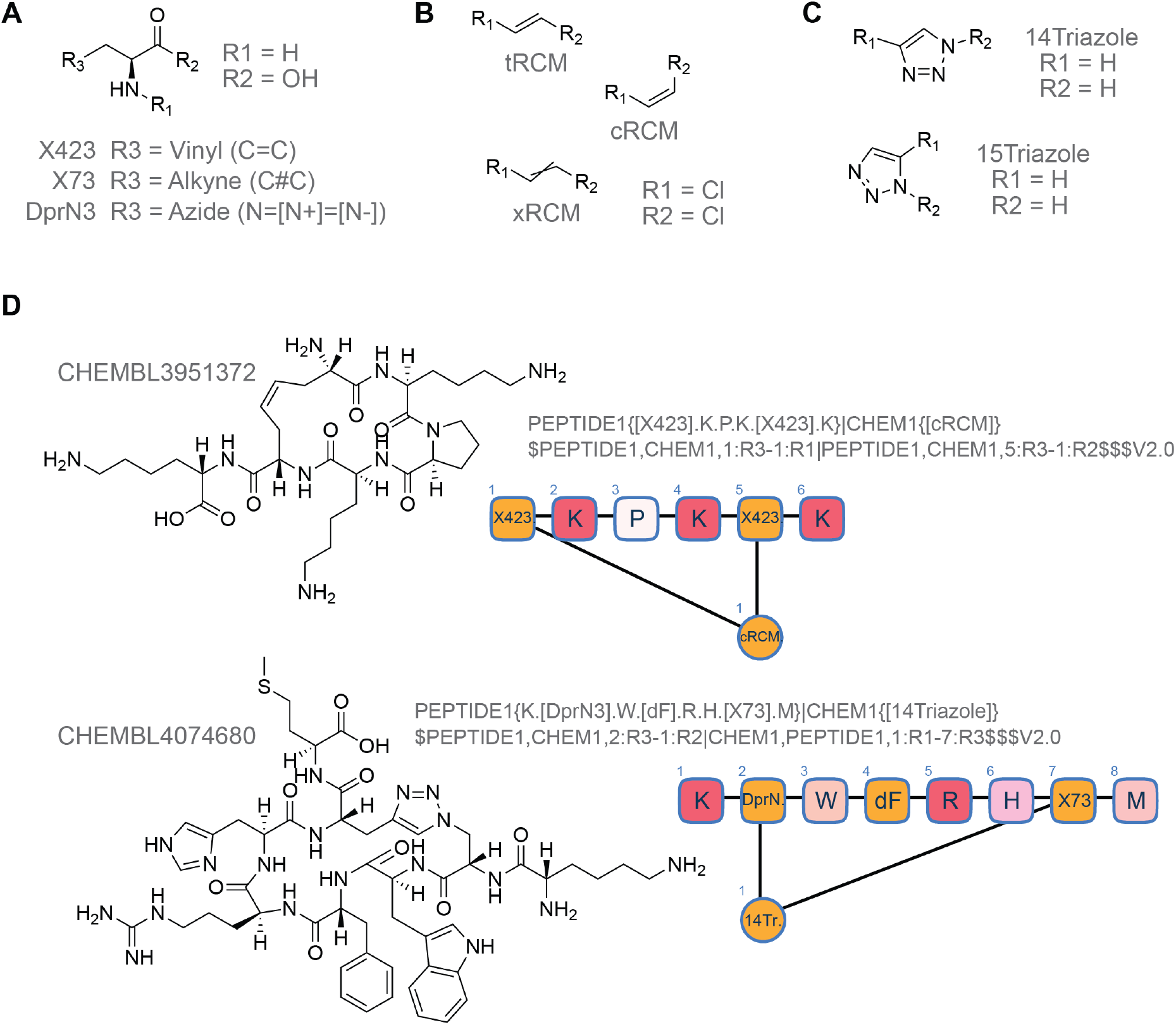
New HELM Rules for Complex Connections in Peptides. a) Examples of Peptide monomers containing the new R Groups. These monomers are used to capture non-natural linkages. Monomer symbols X423 and X73 are taken from the ChEMBL 28 monomer library. Symbol DprN3 was assigned based on structure of Dpr monomer currently available in the CHEMBL 28 monomer library. b) CHEM monomers used for alkene linkages from ring closing metathesis (RCM). c) CHEM monomers used for triazole linkages from Click reactions. d) Examples of added HELM notation for existing ChEMBL compounds generated using the new HELM rules for complex connections.

### Architecture of Tools

PepSeA tools presented herein are designed to be deployed as web services with endpoints that provide Application Programming Interface (API) functionalities for multiple sequence alignment and HELM depiction. Briefly, once the tools are running locally or on a server, the user sends a request with HELM sequences and user-defined parameters to the API for sequence alignment or depiction and the processed result is returned. For sequence alignment, the result includes a detailed breakdown by position of aligned sequences which can be further used for visualization and data analysis. For visualization, the input can be a single HELM string or a list of HELM strings and images (either as Scalable Vector Graphics, SVG, or Portable Network Graphics, PNG) are returned. The deployment of tools as modular API endpoints allows flexible integration with different applications and workflows for data analysis and visualization.

### Improved Depiction and Visual Analysis of Peptide HELM Sequences with PepSeA Viz Tool

Depiction of peptide macrocycles is challenging, especially when they include multiple connections between amino acids. HELM notation offers a suitable alternative to depict biopolymers as sequences while also depicting the structure of monomers (**Figure 2**). Since the inception of the HELM notation, several commercial and open-source depiction tools have been built around the Pistoia Alliance web-based HELM editor.^30^ The PepSeA HELM tool expands the basic HELM depiction capabilities of currently available solutions to enable new visualization features, including a depiction of multiple peptides for sequence analysis in a single visualization (**Figure 2**).

**Figure 2.**
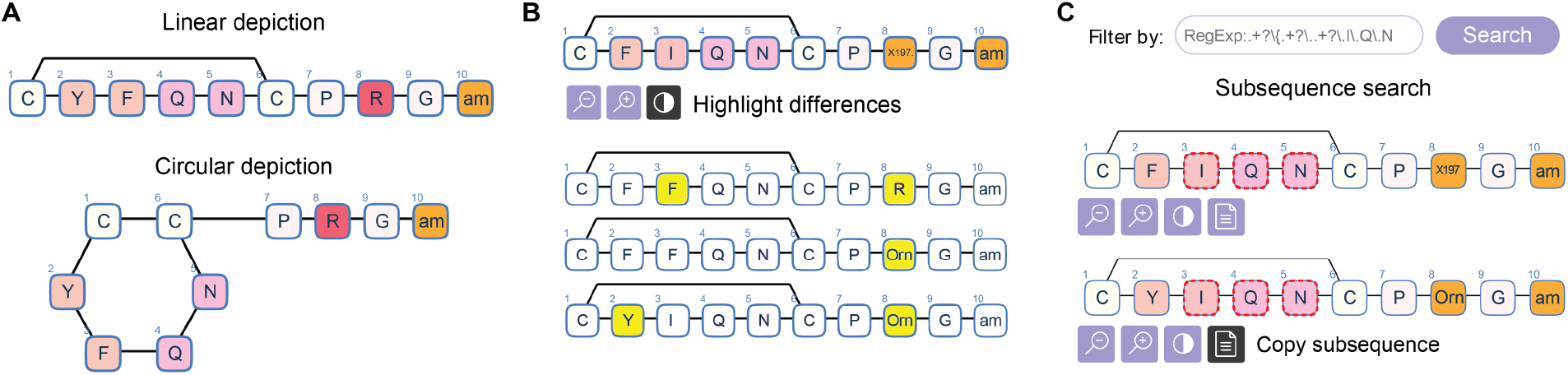
Functionalities of the PepSeA HELM depiction tool. a) Toggling between two different sequence depictions: linear and circular for macrocycles in ChEMBL. b) Changes in sequences of same length can be highlighted using the comparison mode. c) Sequences in selection can be filtered using subsequence search. Example shows sequences with “I.Q.N” HELM subsequence.

The visualization tool allows toggling between linear and cyclic sequence depictions depending on user preferences (**Figure 2**a). Another useful functionality is highlighting the sequence differences for visual identification of peptides with a small number of non-identical residues (**Figure 2**b). This function allows the user to define a reference sequence to be used for positional highlighting by simply selecting it from the visualization. An additional function enables filtering the list of sequences by specific monomers using a substring search (**Figure 2**c). The user can choose a few monomers in a sequence by clicking on a monomer in the depiction and then clicking on the copy button under the sequence. This selection will create a regular expression that can be used for quick filtering of sequences containing the defined pattern. Finally, the PepSeA visualization tool handles a variety of complicated HELM strings, such as multiple macrocyclic peptides that are interconnected (Figure S2). This improvement allows more intuitive inspection of peptides sequences.

### Multiple Sequence Alignment (MSA) of HELM Peptides with Non-Natural Amino Acids

Multiple sequence alignment (MSA) is a well-established bioinformatics technique to analyze sequences of nucleotides or amino acids to identify point mutations, insertions and deletions across sequences.^46^ Although sequence alignment is widely employed to analyze nucleotide and natural amino acid sequences, this method is not directly applicable to peptide sequences containing nonnatural amino acids due to some key limitations. For example, a typical peptide monomer library (such as the CHEMBL 28 HELM monomer library) includes several hundred non-natural amino acids which cannot be accounted for in a conventional substitution matrix built considering evolutionary changes in sequences.^41,47–49^ In addition, most MSA methods can only handle a small number of characters (i.e., 20 one-letter codes of natural amino acids) to distinguish natural nucleotides or amino acids in sequences to align,^50–52^ which significantly limits MSA of peptide sequences with non-natural amino acids.

To overcome these limitations, we developed an MSA strategy in PepSeA, termed MAFFT-ROCS hereafter. This MSA strategy combines a custom substitution matrix generated using ROCS^43^ similarities with MAFFT^35^ sequence alignment that can be directly used for HELM peptide sequences (see Methods). MAFFT-ROCS addresses the abovementioned challenges by generating a custom substitution matrix that takes into account the chemical and structural diversity of monomers in the ChEMBL 28 HELM library and by enabling sequence alignment of HELM peptides with non-natural amino acids using the extended alphabet capability of the MAFFT tool.^35^

To show the usability of the MAFFT-ROCS strategy, we applied it to a subset of the HELM peptides from the ChEMBL database (see Methods for details of subset generation). Since MAFFT offers a variety of different alignment algorithms,^34,35^ different combination of parameters were tested to identify the most applicable configuration for HELM peptides. The datasets were characterized in terms of percentage of sequence similarity and sequence length because these parameters were expected to affect the alignment performance (Table S2). Peptides in the CHEMBL1008221 dataset are of high similarity with an average sequence similiarty of 0.64 and close to a constant length of 12 with a small number of N-terminal insertions. The CHEMBL956457 dataset is of lower similarity (average sequence similarity: 0.49), contains 16-residue long peptides on average, and a small number of circular peptides (5/151). The low-similarity CHEMBL660105 dataset (average sequence similarity: 0.21) consists of same-length peptides (9) with predominantly natural amino-acids and a few C-terminally O-methylated (OMe) residues. Similarity and sequence length distributions are summarized in Table S2 and plotted in Figure S3.

To evaluate the performance of the MAFFT-ROCS strategy, the simulation program ROSE ^53^ was used to generate peptides that resemble the original ChEMBL datasets in terms of sequence similarity and length distributions (Figure S4), and to obtain the true MSA as a reference to evaluate the alignment quality (Table S3). Common measures to evaluate the alignment quality are the Sum of Pairs (SPS) score, i.e., the fraction of aligned pairs, and the Column Score (CS), which is the fraction of columns correctly reproduced in the tested MSA compared to the reference MSA ^54^. The gap opening and extension parameters were varied with values (1, 1.53, 2, 3, 5, 10) and (0, 0.5, 1, 2.5, 5), respectively, and the alignment methods tested were the fast progressive method FFT-NS-2 and the more accurate iterative methods which use either global or local G/L/E-INS-I algorithms in the pairwise alignment stage (referred to as ginsi, linsi, einsi below). From the analysis of simulated datasets, the local alignment method linsi was most accurate for these datasets (Figures S5, S6 and S7, details of analysis are presented in Supporting Information). Also recognized is the sensitivity of the MSA quality to the gap parameters, especially for lower-similarity datasets (details in Supporting Information).

From both MSA MAFFT-ROCS alignment on simulated dataset and the ChEMBL datasets, the linsi method was identified as successful in producing higher quality scores (Figure S5), with default gap parameters not always the best choice as demonstrated on the low-similarity dataset. Given the diversity of potential datasets, users are encouraged to run a similar analysis to identify suitable combination of parameters.

### Visualization and SAR Analysis of Vasopressin dataset enabled by PepSeA tools

To showcase the applicability of the PepSeA tools we analyze potency data of 52 vasopressin analogues (CHEMBL ID: ChEMBL1819839)^33^ that target V1a vasopressin receptor, a receptor that is located in smooth muscle cells and that is involved in vasodilatory hypotension. Using the PepSeA tools offered a more concise way to analyze SAR data previously published (**Figure 3**). Based on the analysis shown in Figure 3b a chemistry team can immediately develop a strategy to (1) keep F in position 3 and replace R with Orn in position 8 and (2) F, Y or Cha in position 2 and Q in position 4 (based on the lasso regression Figure 3d).

**Figure 3.**
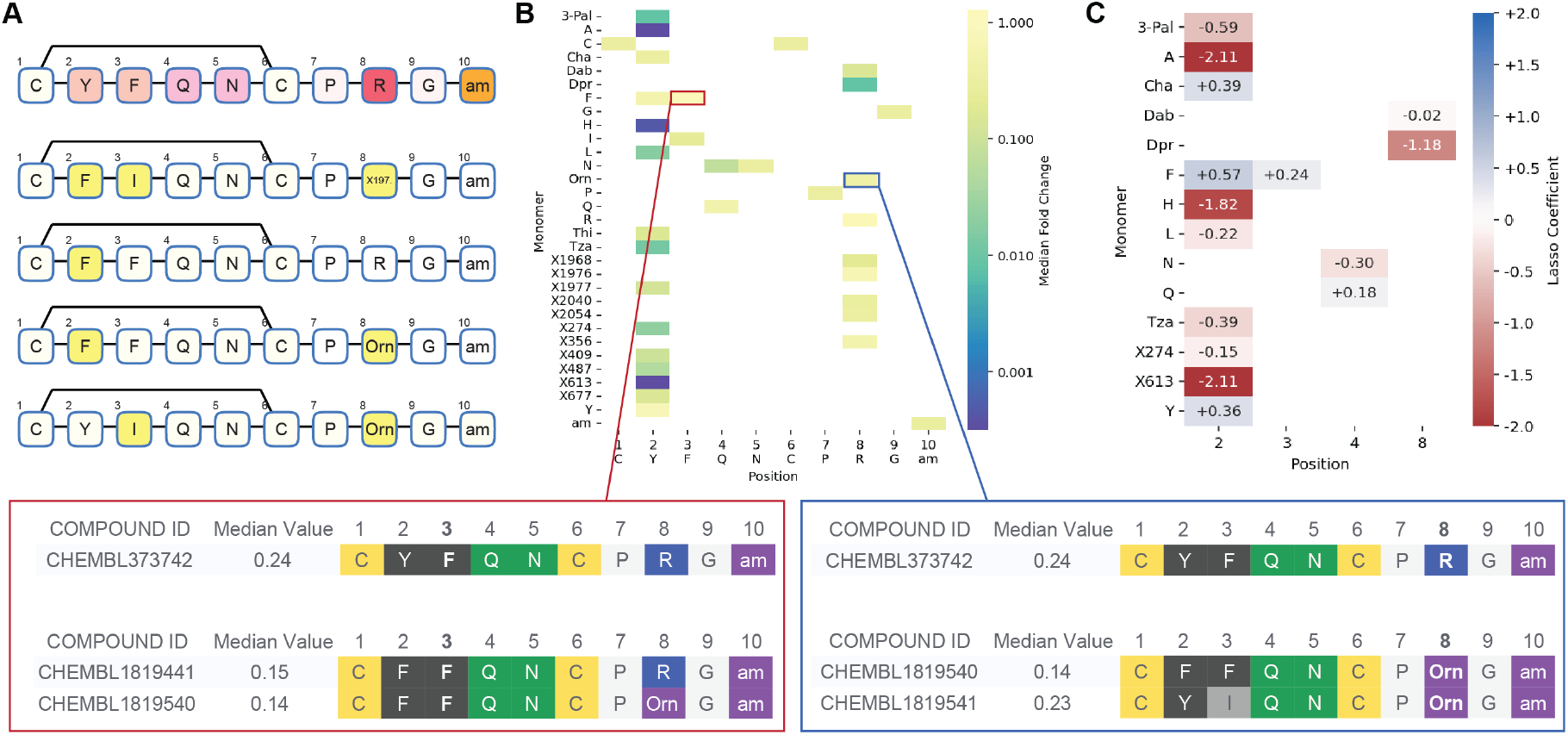
Example of an analysis workflow that combines all the tools to analyze CHEMBL1819839 EC50 potency data. a) Sequence alignment with highlighting of amino acid changes. Vasopressin sequence (CHEMBL373742) is used as a reference. Differences are highlighted in yellow. b) A mutation cliff analysis using aligned sequences. Each box in the heat map represents fold change in EC50 relative to vasopressin reference (ratio of EC50 of contributing peptides and vasopressing reference). Red and blue boxes represent examples of peptides with positive mean fold change as described next. The blue box shows an example of peptides with positive median fold change relative to vasopressin where position 8 is changed from R to Orn. The red box shows peptides where amino acid at position 3 is the same relative to vasopressin, but other positions are different. c) Coefficients of a lasso regression model built using the sequence alignment matrix. Each box in the heat map indicates feature importance (contribution) to overall peptide potency.

The HELM depiction tool allowed a quick inspection of differences in peptide sequences relative to the vasopressin reference sequence (**Figure 3**a). This visualization, coupled with potency data, provided a qualitative way to inspect sequence-activity relationships. A more quantitative analysis can be performed using sequence alignment across all 52 peptide analogues that enables a mutation cliff analysis (**Figure 3**b).^55^ Mutation cliffs are helpful to visualize how changes in sequence affect peptide potency. Each box in the heat map of the mutation cliff plot represents the mean fold change when an amino acid in the reference sequence (vasopressin with CHEMBL ID CHEMBL373742 in this example, shown in the horizontal axis of the plot) is changed to another amino acid at a specific position. In this analysis, the mean fold change is calculated using the average value of the potency taken across all the peptides that have the changed amino acid at that position. For example, a mean fold change of 1.35 results when Arg at position 8 of vasopressin is changed to Orn in peptides CHEMBL1819540 and CHEMBL1819541 (**Figure 3**b, blue box). Another example highlights positive mean fold changes when position 3 is maintained (F in vasopressin), but other changes are made in the sequence in peptides CHEMBL1819441 (F to Y at position 2) and CHEMBL1819540 (R to Orn at position 8 as described before) (**Figure 3**b, red box). This visualization enabled by MAFFT-ROCS sequence alignment with PepSeA provides an intuitive way to explore SAR of peptide sequences. It is important to note that mutation cliff analysis is sensitive to how the underlying positional matrix is built. A poor sequence alignment would result in a skewed mutation cliff matrix which could lead to incorrect assumptions during positional SAR analysis. This highlights the need for a robust sequence alignment scheme that considers non-natural amino acids such as the proposed MAFFT-ROCS strategy.

Another impactful analysis enabled by MAFFT-ROCS sequence alignment is Lasso regression for positional SAR analysis (**Figure 3**c). This analysis provides a quantitative way to analyze how certain amino acids (vertical axis in the plot) at specific positions (the horizontal axis in the plot) drive peptide potency. Lasso regression ranks feature importance and contributions (i.e., amino acids and positions) to potency using regression coefficients that are easy to interpret in the context of the peptide sequence. One potential limitation of this analysis is that it does not consider nonlinear, synergistic relationships between multiple amino acid and position combinations. However, this analysis can be informative for choosing which amino acid and what positions are worth exploring in the next iteration of the peptide design cycle (**Figure 3**c). It is also important to note that, in principle, the rigorous positional matrix obtained with MAFFT-ROCS sequence alignment can be used to generate appropriate input matrices for more sophisticated machine learning algorithms for positional SAR or de novo peptide sequence design.

## CONCLUSION

In this work we described components of PepSeA, a new set of tools for SAR analysis of peptide data, sequence alignment and upgraded depiction of HELM sequences. We have made these tools open-source and freely available to the wider community. To the best of our knowledge, there is currently no off-the-shelf solution for alignment of HELM sequences containing multiple nonnatural amino acids. The PepSeA tools represent an initial attempt to expand cheminformatics capabilities for peptides.

The PepSeA tools are modular and flexible and can be integrated into existing workflows using simple API calls as described earlier. The MAFFT-ROCS strategy at the core of PepSeA sequence alignment addresses the unmet need for sequence-based peptide SAR analysis. The PepSeA sequence alignment of non-natural amino acids enables intuitive ways to get an insight into peptide SAR, such as the mutation cliff analysis. Moreover, the resulting matrix of aligned sequences can be used to build sequence based QSAR models for peptide design. The similarity matrix used to generate the custom substitution matrix can also be used to select amino acids for virtual peptide library designs or guide selection of amino acids at positions deemed important using mutation cliffs analysis.

PepSeA tools can be used to analyze and visualize 2-dimensional data in the form of peptide sequences. A future direction for PepSeA tools could be integration with structure-based peptide design workflows. Integration of the alignment API with a 3D design tool can enable new design insights that leverage information extracted from a set of aligned peptides and 3D interactions with the target protein. Another design enabling possibility would be the integration of the sequencebased workflows enabled by PepSeA with design workflows that make atom-level changes on peptide structures. In this manner, new amino acid substitutions or linkages could be made by modifying peptide structure and subsequently incorporated into an updated HELM sequence.

With the increasing exploration of peptides as therapeutics by discovery teams, enabling rapid SAR visualization and design cycles will become ever more important. PepSeA is a tool that can fill an existing gap in peptide SAR visualization and analysis and enable faster and more productive design–make–test cycles.

## ASSOCIATED CONTENT

### Supporting Information

The following files are available free of charge.

Examples of ChEMBL peptide-like compounds with HELM sequence generated using new rules described in this study (XLSX)

All peptides from ChEMBL (with and without HELM sequences as described in Methods section) and associated activity data (XLSX)

Vassopressin dataset and data used to generate heatmaps in Figure 3 (CSV)

Supplementary Figures and tables, including schematic of ROCS workflow, HELM depiction examples and MSA benchmarking using ChEMBL datasets. (DOCX)

Monomer symbol to ASCII lookup table used for sequence alignment (TXT)

ROCS substitution matrix for ChEMBL monomers (TXT)

## AUTHOR INFORMATION

## Acknowledgments

We thank Jens Christensen, Vincent Antonnuci, Carol A. Rohl and Juan Alvarez for supporting this work. We also thank Sookhee Ha, Nicolas Boyer, Michael Garrigou and Jennifer Hickey for their help and contributions to defining new HELM notation rules. We are immensely grateful to David Prihoda for his help with the artwork.

## Author Contributions

DAB and MG conceived and supervised the study. JB scientifically led and designed the functionalities of PepSeA with significant help from OU, AW and GA. JB and GA devised new rules for HELM notation. JB, OU and AW provided scientific insights for HELM-based sequence alignment. MS designed and led the implementation of the software. PM designed and implemented the MSA strategy using MAFFT. KC and VP developed the HELM visualization. EJ and DB designed integration and guided its implementation by VP. AM and DH implemented and tested different functions of PepSeA and AM performed the benchmarking and comparative analysis of MSA. AG and CC managed the development team and supervised the implementation. JB, DAB, AM, MG, OU and GA wrote the manuscript. All authors read and approved the final version of the manuscript.

## Funding Sources

This work was supported by Merck Sharp & Dohme Corp., a subsidiary of Merck & Co., Inc., Kenilworth, NJ, USA.

## Notes

All code for this publication is available in the following GitHub repository: https://github.com/Merck/PepSeA.

## Competing Interests

The authors declare no conflict of interest.

## Notes

https://github.com/Merck/PepSeA

### Competing Interest Statement

The authors have declared no competing interest.

