## Supplementary Figures for "PepSeA: Peptide Sequence Alignment and Visualization Tools to Enable Lead Optimization"

**Schematic of ROCS workflow**

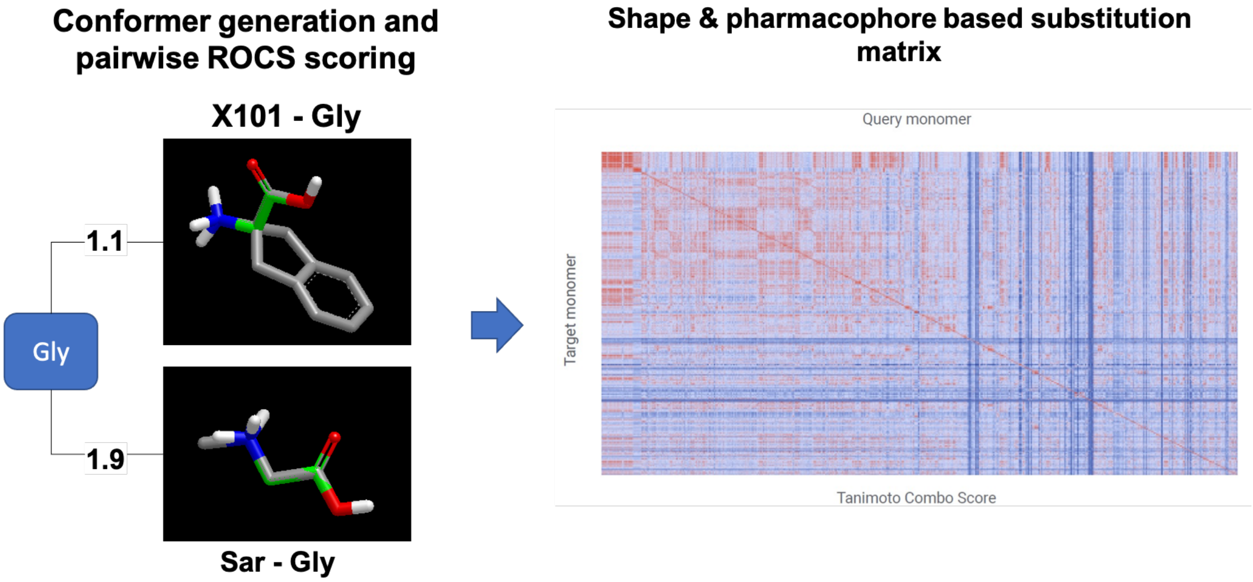


**Figure S1.** Schematic showing the steps to generate ROCS substitution matrix. The custom substation matrix is used for alignment of HELM peptides with non-natural amino acids using MAFFT. On the left side, the example shows ROCS scoring for X101 and Sar from CHEMBL 28 monomer database against Gly. Once scoring is performed for each pair of monomers in monomer library, the best Tanimoto Combo score is selected and rescaled to [-10,10] range to generate the substitution matrix, shown on the right side.

### Example of HELM Depictions

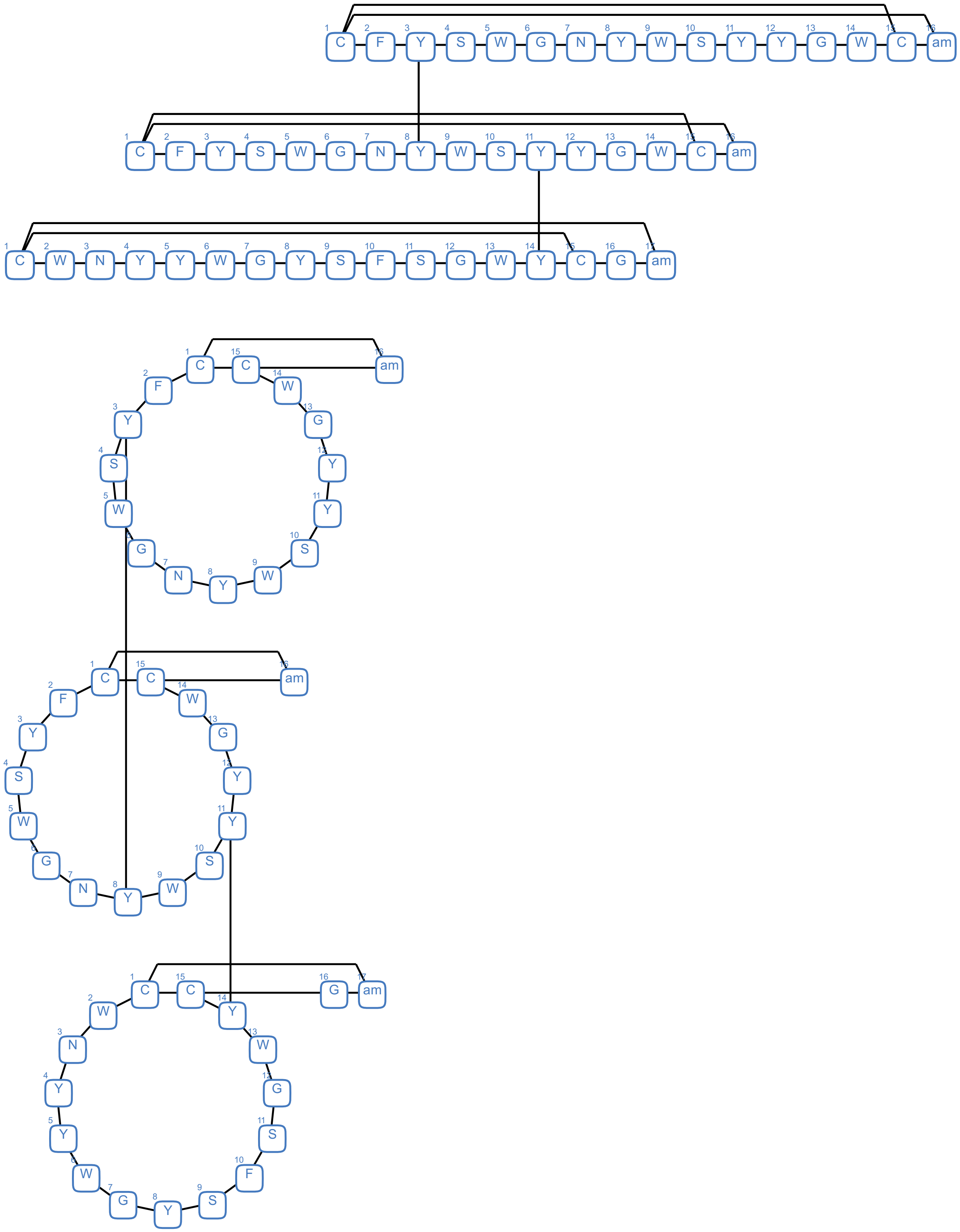


**Figure S2.** Example of improved HELM depiction for interconnected macrocycles.

### MSA Benchmarking with ChEMBL datasets

To show the applicability of the multiple sequence alignment using the ROCS-scoring matrix, we apply it to the three ChEMBL datasets containing non-natural amino-acids and at least 150 HELM sequences. The following datasets, referred to as dataset 0, 1 and 2, respectively, were chosen: target ID: CHEMBL2611, assay ID: CHEMBL1008221, with nanomolar inhibitor peptides against human recombinant furin,[1] target ID: 2632, assay ID: CHEMBL660105, with $IC_{50}$ binding affinity data for peptides targeting the T-cell major histocompatibility complex (MHC),[2] target ID: 5966, assay ID: 660105, with dissociation constants for peptides targeting the FcRn receptor, the key regulatory protein for IgG gamma-globulin homeostasis.[3] The sequences of the peptides and their corresponding experimental activities are listed in Table S1.

Table S1. ChEMBL datasets used in MSA-Benchmarking

Dataset 0

| Peptide Sequence | $K_i$ / nM |
|---|---|
| TPRERRRKKRG[NH2] | 100 |
| TPRARRRKKR[Nle][NH2] | 81 |
| TPRARRRKKR[X202][NH2] | 65 |
| TPRARRRKKRR[NH2] | 100 |
| TPRARRRKKRK[NH2] | 100 |
| TPRARRRKKRH[NH2] | 100 |
| TPRARRRKKRE[NH2] | 100 |
| TPRARRRKKRD[NH2] | 56 |
| TPRARRRKKRQ[NH2] | 47 |
| TPRARRRKKRN[NH2] | 45 |
| TPRARRRKKRY[NH2] | 53 |

| | |
|---|---|
| TPRARRRKKRC[NH2] | 100 |
| TPRARRRKKRT[NH2] | 23 |
| TPRARRRKKRS[NH2] | 100 |
| TPRARRRKKRM[NH2] | 100 |
| TPRARRRKKRW[NH2] | 59 |
| TPRARRRKKRF[NH2] | 38 |
| TPRARRRKKRP[NH2] | 100 |
| TPRARRRKKRI[NH2] | 30 |
| TPRARRRKKRL[NH2] | 100 |
| TPRARRRKKRV[NH2] | 80 |
| TPRARRRKKRA[NH2] | 100 |
| TPR[dK]RRRKKRG[NH2] | 100 |
| [alT]PR[dgE]RRRKKRG[NH2] | 100 |
| TPR[Cit]RRRKKRG[NH2] | 100 |
| TPR[Acc5]RRRKKRG[NH2] | 100 |
| TPRRRRRKKRG[NH2] | 100 |
| TPRKRRRKKRG[NH2] | 100 |
| TPRHRRRKKRG[NH2] | 100 |
| TPRDRRRKKRG[NH2] | 100 |
| TPRQRRRKKRG[NH2] | 100 |
| TPRNRRRKKRG[NH2] | 100 |
| TPRYRRRKKRG[NH2] | 100 |
| TPRCRRRKKRG[NH2] | 100 |
| TPRTRRRKKRG[NH2] | 100 |
| TPRSRRRKKRG[NH2] | 100 |
| TPRMRRRKKRG[NH2] | 100 |
| TPRWRRRKKRG[NH2] | 100 |
| TPRFRRRKKRG[NH2] | 100 |
| TPRPRRRKKRG[NH2] | 100 |
| TPRIRRRKKRG[NH2] | 100 |

| | |
|---|---|
| TPRLRRRKKRG[NH2] | 100 |
| TPRVRRRKKRG[NH2] | 100 |
| TPRGRRRKKRG[NH2] | 100 |
| TPRERRRKKAG[NH2] | 100 |
| TPRERRRKARG[NH2] | 100 |
| TPRERRRAKRG[NH2] | 100 |
| TPRERRAKKRG[NH2] | 100 |
| TPRERARKKRG[NH2] | 100 |
| TPREARRKKRG[NH2] | 100 |
| TPRARRRKKRG[NH2] | 57 |
| TPAERRRKKRG[NH2] | 100 |
| TARERRRKKRG[NH2] | 100 |
| [dA]PRERRRKKRG[NH2] | 100 |
| TPQRARRRKKRT | 33 |
| TPQRARRRKKRY | 47 |
| TPQRARRRKKRF | 38 |
| TPQRARRRKKRW | 34 |
| TPRERRRKKR | 100 |
| TPREAAAKKR | 100 |
| TPAAARRKKR | 100 |
| AAAERRRKKR | 100 |
| TPRERRRKAA | 100 |
| TPRERRRAAR | 100 |
| TPRERRAAKR | 100 |
| TPRERAAKKR | 100 |
| TPREAARKKR | 100 |
| TPRAARRKKR | 100 |
| TPAARRRKKR | 100 |
| TAAERRRKKR | 100 |
| AARERRRKKR | 100 |

| | |
|---|---|
| TPRERRRKKA | 100 |
| TPRERRRKAR | 100 |
| TPRERRRAKR | 100 |
| TPRERRAKKR | 100 |
| TPRERARKKR | 100 |
| TPREARRKKR | 100 |
| TPRARRRKKR | 138 |
| TPAERRRKKR | 100 |
| TARERRRKKR | 100 |
| APRERRRKKR | 100 |
| AARERRRKKRG | 100 |
| TPREAAAKKR[Sar] | 100 |
| TPAAARRKKR[Sar] | 100 |
| AAAERRRKKR[Sar] | 100 |
| TPRERRRKAA[Sar] | 100 |
| TPRERRRAAR[Sar] | 100 |
| TPRERRAAKR[Sar] | 100 |
| TPRERAAKKR[Sar] | 100 |
| TPREAARKKR[Sar] | 100 |
| TPRAARRKKR[Sar] | 100 |
| TPAARRRKKR[Sar] | 100 |
| TAAERRRKKR[Sar] | 100 |
| AARERRRKKR[Sar] | 100 |
| TPRERRRKKA[Sar] | 100 |
| TPRERRRKAR[Sar] | 100 |
| TPRERRRAKR[Sar] | 100 |
| TPRERRAKKR[Sar] | 100 |
| TPRERARKKR[Sar] | 100 |
| TPREARRKKR[Sar] | 100 |
| TPRARRRKKR[Sar] | 100 |

| | |
|---|---|
| TPAERRRKKR[Sar] | 100 |
| TARERRRKKR[Sar] | 100 |
| APRERRRKKR[Sar] | 100 |
| TPRERRRKKR[Sar] | 100 |
| TPREAAAKKRV | 100 |
| TPAAARRKKRV | 100 |
| AAAERRRKKRV | 100 |
| TPRERRRKAAV | 100 |
| TPRERRRAARV | 100 |
| TPRERRAAKRV | 100 |
| TPRERAAKKRV | 100 |
| TPREAARKKRV | 100 |
| TPRAARRKKRV | 100 |
| TPAARRRKKRV | 100 |
| TAAERRRKKRV | 100 |
| AARERRRKKRV | 100 |
| TPRERRRKKAV | 100 |
| TPRERRRKARV | 100 |
| TPRERRRAKRV | 100 |
| TPRERRAKKRV | 100 |
| TPRERARKKRV | 100 |
| TPREARRKKRV | 100 |
| TPAERRRKKRV | 100 |
| TARERRRKKRV | 100 |
| APRERRRKKRV | 100 |
| TPRERRRKKRV | 100 |
| TPREAAAKKRG | 100 |
| TPAAARRKKRG | 100 |
| AAAERRRKKRG | 100 |
| TPRERRRKAAG | 100 |

| | |
|---|---|
| TPRERRRAARG | 100 |
| TPRERRAAKRG | 100 |
| TPRERAAKKRG | 100 |
| TPREAARKKRG | 100 |
| TPRAARRKKRG | 100 |
| TPAARRRKKRG | 100 |
| TAAERRRKKRG | 100 |
| GGGGGGTPQRARRRKKRW | 88 |
| TPQRARRRKKRG | 100 |
| TPQRARRRKKR[dK] | 100 |
| TPQRARRRKKR[Nle] | 100 |
| TPQRARRRKKR[X202] | 93 |
| TPQRARRRKKR[Acc5] | 100 |
| TPQRARRRKKRR | 100 |
| TPQRARRRKKRK | 100 |
| TPQRARRRKKRH | 100 |
| TPQRARRRKKRE | 103 |
| TPQRARRRKKRD | 50 |
| TPQRARRRKKRQ | 100 |
| TPQRARRRKKRN | 100 |
| TPQRARRRKKRC | 100 |
| TPQRARRRKKRS | 100 |
| TPQRARRRKKRM | 58 |
| TPQRARRRKKRP | 100 |
| TPQRARRRKKRI | 100 |
| TPQRARRRKKRL | 100 |
| TPQRARRRKKRV | 48 |
| TPQRARRRKKRA | 115 |
| TPQRARRRKKR[dR] | 100 |
| TPQR[Acc5]RRRKK[Spg]G | 100 |

| | |
|---|---|
| TPQR[orAbu]RRRKKRG | 100 |
| TPQR[Cha]RRRKKRG | 100 |
| TPQR[Nle]RRRKKRG | 100 |
| TPQR[X202]RRRKKRG | 100 |
| TPQR[dR]RRRKKRG | 100 |
| TPQR[dK]RRRKKRG | 100 |
| TPQR[dgE]RRRKKRG | 100 |
| TPQR[Cit]RRRKKRG | 100 |
| TPQR[Acc5]RRRKKRG | 100 |
| TPQRYRRRKKRG | 100 |
| TPQRWRRRKKRG | 100 |
| TPQRVRRRKKRG | 100 |
| TPQRTRRRKKRG | 100 |
| TPQRSRRRKKRG | 100 |
| TPQRRRRRKKRG | 100 |
| TPQRQRRRKKRG | 100 |
| TPQRPRRRKKRG | 100 |
| TPQRNRRRKKRG | 100 |
| TPQRMRRRKKRG | 100 |
| TPQRLRRRKKRG | 100 |
| TPQRKRRRKKRG | 100 |
| TPQRIRRRKKRG | 100 |
| TPQRHRRRKKRG | 100 |
| TPQRGRRRKKRG | 100 |
| TPQRFRRRKKRG | 100 |
| TPQRERRRKKRG | 100 |
| TPQRDRRRKKRG | 100 |
| TPQRCRRRKKRG | 100 |
| TPQRARRRKK[Cit]G | 100 |
| TPQRARRRKK[dK]G | 100 |

| | |
|---|---|
| TPQRARRRKK[dR]G | 100 |
| GGGTPRARRRKKRT | 39 |
| GGGGGGTPRARRRKKRT | 66 |
| GGGTPQRARRRKKRW | 67 |
| GAGAGATPRARRRKKRY | 57 |
| GGGGGGTPRARRRKKRY | 54 |
| GGGTPRARRRKKRY | 42 |
| GAGAGATPRARRRKKRF | 56 |
| GGGGGGTPRARRRKKRF | 45 |
| GGGTPRARRRKKRF | 67 |
| GAGAGATPRARRRKKRT | 47 |
| TPRERRRKKAG | 100 |
| TPRERRRKKRG | 100 |
| TPRERRRKARG | 100 |
| TPRERRRAKRG | 100 |
| TPRERRAKKRG | 100 |
| TPRERARKKRG | 100 |
| TPREARRKKRG | 100 |
| TPRARRRKKRG | 57 |
| TPAERRRKKRG | 100 |
| TARERRRKKRG | 100 |
| APRERRRKKRG | 100 |
| TPRARRRKKRV | 80 |
| TPRARRRKKRA | 100 |
| TPRARRRKKRF | 38 |
| TPRARRRKKRP | 100 |
| TPRARRRKKRI | 30 |
| TPRARRRKKRL | 100 |
| TPRARRRKKRW | 59 |
| TPRARRRKKR[Nle] | 81 |

| | |
|---|---|
| TPRARRRKKR[X202] | 65 |
| TPRARRRKKRR | 100 |
| TPRARRRKKRK | 100 |
| TPRARRRKKRH | 100 |
| TPRARRRKKRE | 100 |
| TPRARRRKKRD | 56 |
| TPRARRRKKRQ | 47 |
| TPRARRRKKRN | 45 |
| TPRARRRKKRY | 53 |
| TPRARRRKKRC | 100 |
| TPRARRRKKRT | 23 |
| TPRARRRKKRS | 100 |
| TPRARRRKKRM | 100 |
| TPRCRRRKKRG | 100 |
| TPRQRRRKKRG | 100 |
| TPRPRRRKKRG | 100 |
| TPRNRRRKKRG | 100 |
| TPRMRRRKKRG | 100 |
| TPRLRRRKKRG | 100 |
| TPRKRRRKKRG | 100 |
| TPRIRRRKKRG | 100 |
| TPRHRRRKKRG | 100 |
| TPRGRRRKKRG | 100 |
| TPRFRRRKKRG | 100 |
| TPRDRRRKKRG | 100 |
| TPR[dK]RRRKKRG | 100 |
| TPR[dgE]RRRKKRG | 100 |
| TPR[Cit]RRRKKRG | 100 |
| TPR[Acc5]RRRKKRG | 100 |
| TPRYRRRKKRG | 100 |

| | |
|---|---|
| TPRWRRRKKRG | 100 |
| TPRVRRRKKRG | 100 |
| TPRTRRRKKRG | 100 |
| TPRSRRRKKRG | 100 |
| TPRRRRRKKRG | 100 |

### Dataset 1

| Peptide Sequence | $IC_{50}$ / nM |
|---|---|
| GIGILTVIL | 1000 |
| HLYSHPIIL | 73.96 |
| GTLGIVCPI | 193.2 |
| GTLVALVGL | 4549.88 |
| LQTTIHDII | 3155 |
| DPKVKQWPL | 666.81 |
| GLGQVPLIV | 500.03 |
| KLPQLCTEL | 328.1 |
| HLAVIGALL | 100 |
| ALAKAAAAM | 39.99 |
| MLDLQPETT | 462.38 |
| YTDQVPFSV | 85.9 |
| VVMGTLVAL | 66.99 |
| LLFGYPVYV | 13 |
| FLDQVPFSV | 2.198 |
| IISCTCPTV | 263.03 |
| YLYPGPVTV | 8.892 |
| LLSCLGCKI | 3572.73 |
| ILKEPVHGV | 11.99 |
| WLDQVPFSV | 11.51 |
| GILTVILGV | 4.498 |

| | |
|---|---|
| LLVVMGTLV | 1352.07 |
| ILTVILGVL | 381.07 |
| ITMQVPFSV | 39.99 |
| GLACHQLCA | 416.87 |
| ALAKAAAAL | 308.32 |
| PLLPIFFCL | 159.96 |
| SLDDYNHLV | 26 |
| MLGTHTMEV | 14.29 |
| YLFPGPVTA | 3.199 |
| YLMPGPVTA | 4.295 |
| WILRGTSFV | 277.97 |
| YMIMVKCWM | 217.27 |
| YVITTQHWL | 103.99 |
| LLAQFTSAI | 50 |
| IIDQVPFSV | 39.99 |
| YAIDLPVSV | 16 |
| ILWQVPFSV | 1.698 |
| AAAKAAAAV | 399.94 |
| KLHLYSHPI | 29.38 |
| NLSWLSLDV | 229.61 |
| FVTWHRYHL | 944.06 |
| ILHNGAYSL | 74.64 |
| RLMKQDFSV | 45.5 |
| NLYVSLLLL | 76.91 |
| QLFEDNYAL | 17.22 |
| SLHVGTQCA | 1438.8 |
| ALAKAAAAA | 112.98 |
| KTWGQYWQV | 11.09 |
| YLDQVPFSV | 2.301 |
| VVLGVVFGI | 14.29 |

| | |
|---|---|
| QVMSLHNLV | 676.08 |
| LLGCAANWI | 5000.35 |
| QLFHLCLII | 130.02 |
| ALPYWNFAT | 1352.07 |
| VLHSFTDAI | 416.87 |
| ALCRWGLLL | 100 |
| ILMQVPFSV | 7.499 |
| TLGIVCPIC | 153.11 |
| LLCLIFLLV | 100.93 |
| SLNFMGYVI | 1315.22 |
| SIISAVVGI | 69.34 |
| GLYSSTVPV | 26.52 |
| ILSQVPFSV | 20 |
| HLYQGCQVV | 147.23 |
| ILLLCLIF[L_OMe] | 142.89 |
| FAFRDLCI[V_OMe] | 130.02 |
| VLLDYQGML | 46.99 |
| NMVPFFPPV | 3.999 |
| TVILGVLLL | 847.23 |
| AAGIGILTV | 262.42 |
| YMLDLQPET | 48.98 |
| WLSLLVPFV | 8.954 |
| ITAQVPFSV | 95.5 |
| ITYQVPFSV | 33.11 |
| ILFQVPFSV | 2 |
| LTVILGVLL | 2630.27 |
| AIAKAAAAV | 666.81 |
| MLLAVLYCL | 333.43 |
| FTDQVPFSV | 61.38 |
| KIFGSLAFL | 33.27 |

| | |
|---|---|
| YLEPGPVTL | 87.5 |
| GLSRYVARL | 56.49 |
| ILSPFMPLL | 44.98 |
| SVYDFFVWL | 35.97 |
| TLDSQVMSL | 161.06 |
| ALIHHNTHL | 238.23 |
| VCMTVDSLV | 7144.96 |
| AVAKAAAAV | 319.89 |
| MMWYWGPSL | 11.99 |
| YLWPGPVTA | 3.199 |
| VLQAGFFLL | 207.97 |
| ALMPLYACI | 10 |
| LLWFHISCL | 207.97 |
| YLYPGPVTA | 16.9 |
| LLSSNLSWL | 454.99 |
| ITFQVPFSV | 66.22 |
| ALAKAAAAV | 381.07 |
| LLLCLIFLL | 26 |
| RLLQETELV | 20.8 |
| WTDQVPFSV | 716.14 |
| YMNGTMSQV | 39.99 |
| FLLTRILTI | 7.096 |
| ITSQVPFSV | 636.8 |
| LMAVVLASL | 111.17 |
| IMDQVPFSV | 19.1 |
| YLEPGPVTA | 214.78 |
| VTWHRYHLL | 161.06 |
| ALAKAAAAI | 615.18 |
| SLYADSPSV | 21.98 |
| YLAPGPVTA | 9.29 |

| | |
|---|---|
| TTAEEAAGI | 4168.69 |
| LLAVGATKV | 333.43 |
| FLEPGPVT[A_OMe] | 126.47 |
| YLSPGPVTV | 22.8 |
| LLMGTLGIV | 7.998 |
| HLESLFTAV | 5000.35 |
| NLQSLTNLL | 1000 |
| WLEPGPVTA | 827.94 |
| YLEPGPVTI | 65.01 |
| YLEPGPVTV | 45.5 |
| VMGTLVALV | 27.99 |
| SAANDPIFV | 4549.88 |
| NLGNLNVSI | 76.03 |
| YLFPGPVTV | 5.794 |
| VALVGLFVL | 7144.96 |
| LIGNESFAL | 384.59 |
| VILGVLLLI | 164.06 |
| FLCKQYLN[L_OMe] | 133.35 |
| TLLVVMGTL | 2630.27 |
| AMFQDPQER | 1819.7 |
| CLTSTVQLV | 147.23 |
| VLIQRNPQL | 22.7 |
| TLHEYMLDL | 187.93 |
| ALVGLFVLL | 26 |
| ITWQVPFSV | 34.43 |
| ALMDKSLHV | 16.98 |
| FLGGTPVCL | 238.23 |
| ILDEAYVMA | 238.23 |
| YLWPGPVTV | 7.499 |
| FVWLHYYSV | 15 |

| | |
|---|---|
| YLMPGPVTV | 11.69 |
| YLAPGPVTV | 15.21 |
| ILAQVPFSV | 11.51 |
| FLLSLGIHL | 8.851 |
| YLSPGPVTA | 41.4 |
| GLLGWSPQA | 5.794 |
| ILYQVPFSV | 4.898 |
| ILDQVPFSV | 3.304 |
| ITDQVPFSV | 112.98 |

### Dataset 2

| Peptide Sequence | $K_d$ / nM |
|---|---|
| QRFCTGHFGGLYPCNGP | 5700 |
| RFCTGHFGGLYPC | 10000 |
| QRFCTGHFGGLYPCN[NH2] | 6100 |
| QRFCTGHFGGLYPCNG | 4600 |
| FCTGHFGGLYPCNGP | 11000 |
| RFCTGHFGGLYPCNGP | 2900 |
| TGHFGGLYP | 250000 |
| CTGHFGGLYPC | 20000 |
| QRFCTGHFGGLYPC | 4200 |
| CTGHFGGLYPCNGP | 34000 |
| QRFCTGHFGGLYPCNGP | 5100 |
| NSFCRGRPGHFGGCYLF | 9400 |
| PSYCIEGHIDGIYCFNA | 8800 |
| KIICSPGHFGGMYCQGK | 22000 |
| GGGCV[alT]GHFGGIYCNYQ | 5200 |
| QRFCTGHF[dA][dA]LYPCNGP | 22000 |
| QRFCTGHFG[dA]LYPCNGP | 12000 |

| | |
|---|---|
| QRFCTGHF[dA]GLYPCNGP | 10000 |
| QRFCT[dA]HFGGLYPCNGP | 250000 |
| QRF[Pen]TGHFG[Sar][NMeL]YPCNGP | 46 |
| RF[Pen]TGHFG[Sar]LYPC | 58 |
| RF[Pen]TGHFGG[NMeL]YPC | 59 |
| QRF[Pen]TGHFGGL[NMeY]PCNGP | 92000 |
| QRF[Pen]TGHFGG[NMeL]YPCNGP | 86 |
| QRF[Pen]TGHFG[Sar]LYPCNGP | 110 |
| QRF[Pen]TGHF[Sar]GLYPCNGP | 2000 |
| QRF[Pen]TGH[NMeF]GGLYPCNGP | 250000 |
| QRF[Pen]T[Sar]HFGGLYPCNGP | 88000 |
| QRFC[NMeA]GHFGGLYPCNGP | 18000 |
| RF[Pen]TGHFGGLYPC | 170 |
| F[Pen]TGHFGGLYPC | 310 |
| QRF[hC]TGHFGGLYP[Pen]NGP | 2100 |
| QRF[Pen]TGHFGGLYP[hC]NGP | 310 |
| QRF[Pen]TGHFGGLYP[Pen]NGP | 370 |
| QRFCTGHFGGLYP[Pen]NGP | 2700 |
| QRF[Pen]TGHFGGLYPCNGP | 250 |
| QRF[dC]TGHFGGLYP[dC]NGP | 200000 |
| QRFCTGHFGGLYP[dC]NGP | 31000 |
| QRF[dC]TGHFGGLYPCNGP | 150000 |
| QRF[dhC]TGHFGGLYPCNGP | 3800 |
| QRFCTGHFGGLYP[dhC]NGP | 3900 |
| QRFCTGHFGGLYPCAGP | 8000 |
| QRFCTGHFGGLYACNGP | 14000 |
| QRFCTGHFGGLAPCNGP | 250000 |
| QRFCTGHFGGAYPCNGP | 26000 |
| QRFCTGHFGALYPCNGP | 120000 |
| QRFCTGHFAGLYPCNGP | 230000 |

| | |
|---|---|
| QRFCTGHAGGLYPCNGP | 250000 |
| QRFCTGAFGGLYPCNGP | 250000 |
| QRFCTAHFGGLYPCNGP | 250000 |
| QRFCAGHFGGLYPCNGP | 4900 |
| QRACTGHFGGLYPCNGP | 28000 |
| QAFCTGHFGGLYPCNGP | 7700 |
| QRF[Pen]TGHFG[dP]LYPCNGP | 230 |
| QRF[dPen]TGHFGGLY[dP][dC]NGP | 250000 |
| QRF[dPen]TGHFGGL[dY]P[dC]NGP | 250000 |
| QRF[dPen]TGH[dF]GG[dL]YP[dC]NGP | 250000 |
| QRF[dPen]TGH[dF]GGLYP[dC]NGP | 250000 |
| QRF[dPen][dalloT]GHFGGLYP[dC]NGP | 250000 |
| QR[dF][dPen]TGHFGGLYP[dC]NGP | 1800 |
| Q[dR]F[dPen]TGHFGGLYP[dC]NGP | 280 |
| QRF[Pen]TGHF[dA][Sar]LYPCNGP | 190 |
| QRF[Pen]TGHF[dP]PLYPCNGP | 100000 |
| QRF[Pen]TGHF[dF]GPYPCNGP | 18300 |
| QRF[Pen]TGHF[dA]G[NMeL]YPCNGP | 230 |
| QRF[Pen]TGHF[dF]G[NMeL]YPCNGP | 240 |
| QRF[Pen]TGHF[dP][dP]LYPCNGP | 3300 |
| QRF[Pen]TGHF[dF][dA]LYPCNGP | 410 |
| QRF[Pen]TGHF[dA][dP]LYPCNGP | 430 |
| QRF[Pen]TGHF[dA][dA]LYPCNGP | 450 |
| QRF[Pen]TGHF[dA]GLYPCNGP | 490 |
| QRF[Pen]TGHFG[dA]LYPCNGP | 480 |
| QRF[Pen]TGHF[Aib]GLYPCNGP | 5200 |
| QRF[Pen]TGHF[dY]GLYPCNGP | 320 |
| QRF[Pen]TGHF[dF]GLYPCNGP | 510 |
| QRF[Pen]TGHF[dalI]GLYPCNGP | 2600 |
| QRF[Pen]TGHF[dH]GLYPCNGP | 410 |

| | |
|---|---|
| QRF[Pen]TGHF[dR]GLYPCNGP | 310 |
| QRF[Pen]TGHF[dP]GLYPCNGP | 790 |
| QRF[Pen]TGHF[dAspO]GLYPCNGP | 580 |
| QRF[Pen]TGHFG[Aib]LYPCNGP | 480 |
| QRF[Pen]TGHFG[dY]LYPCNGP | 1500 |
| QRF[Pen]TGHFG[dF]LYPCNGP | 1700 |
| QRF[Pen]TGHFG[dalI]LYPCNGP | 2200 |
| QRF[Pen]TGHFG[dH]LYPCNGP | 2000 |
| QRF[Pen]TGHFG[dR]LYPCNGP | 830 |
| QRF[dPen]TGH[Tic]GGLYPCNGP | 250000 |
| QRF[Pen]TGH[Phg]GGLYPCNGP | 250000 |
| QRF[Pen]TGHWGGLYPCNGP | 2700 |
| QRF[Pen]TGH[Cha]GGLYPCNGP | 4500 |
| QRF[Pen]TGH[hF]GGLYPCNGP | 7800 |
| QRF[dPen]TGH[Phe(4-Me)]GGLYP[dC]NGP | 200 |
| QRF[dPen]TGH[Phe3Me]GGLYP[dC]NGP | 670 |
| QRF[dPen]TGH[Phe4F43]GGLYP[dC]NGP | 200 |
| QRF[dPen]TGH[2Nal]GGLYP[dC]NGP | 11000 |
| QRF[dPen]TGH[12Nal]GGLYP[dC]NGP | 2200 |
| QRF[dPen]TGH[Phe4F3]GGLYP[dC]NGP | 84000 |
| QRF[dPen]TGH[3Pal]GGLYP[dC]NGP | 19000 |
| QRF[dPen]TGH[2Pal]GGLYP[dC]NGP | 1200 |
| QRF[dPen]TGH[PheF5]GGLYP[dC]NGP | 70000 |
| QRF[dPen]TGH[Acc672]GGLYP[dC]NGP | 18000 |
| QRF[dPen]TGH[Phe(4-NH2)]GGLYP[dC]NGP | 1000 |
| QRFCTGHFGGLFPCNGP | 230000 |
| QRF[Pen]TGHFGGL[Phe4F]PCNGP | 2200 |
| QRF[Pen]TGHFGGL[Acc6500]PCNGP | 290000 |
| QRF[Pen]TGHFGGL[Phe4F3]PCNGP | 180000 |
| QRF[Pen]TGHFGGL[3Pal]PCNGP | 34000 |

| | |
|---|---|
| QRF[Pen]TGHFGGL[2Pal]PCNGP | 120000 |
| QRF[Pen]TGHFGGL[PheF5]PCNGP | 72000 |
| QRF[Pen]TGHFGGL[Acc672]PCNGP | 31000 |
| QRF[Pen]TGHFGGL[Phe(4-NH2)]PCNGP | 34000 |
| QRF[Pen]TGH[hF]G[Sar][NMeL]YPC | 230000 |
| QRF[Pen]TGH[Phe4tbutyl]GGLYPC | 230000 |
| QRF[Pen]TGH[Acc614]GGLYPC | 230000 |
| QRF[Pen]TGH[Dip]GGLYPC | 230000 |
| QRF[Pen]TGH[Phe(4-Cl)]GGLYPC | 230000 |
| QRF[Pen]TGH[Phe(3-Cl)]GGLYPC | 230000 |
| QRF[Pen]TGH[Phe2Cl]GGLYPC | 230000 |
| RF[Pen]TGAFG[Sar][NMeL]YPC | 9900 |
| RF[Pen]TGFFG[Sar][NMeL]YPC | 3300 |
| RF[Pen]TG[Acc6355]FG[Sar][NMeL]YPC | 5200 |
| RF[Pen]TG[dAhe3]FG[Sar][NMeL]YPC | 660 |
| RF[Pen]TG[Acc6015]FG[Sar][NMeL]YPC | 8400 |
| RF[Pen]TG[Phe(4-NH2)]FG[Sar][NMeL]YPC | 220 |
| RF[Pen]TG[Acc662]FG[Sar][NMeL]YPC | 74 |
| RF[Pen]TGRFG[Sar][NMeL]YPC | 500 |
| RF[Pen]TGKFG[Sar][NMeL]YPC | 1300 |
| RF[Pen]TG[Orn]FG[Sar][NMeL]YPC | 1300 |
| RF[Pen]TG[Dab]FG[Sar][NMeL]YPC | 1200 |
| RF[Pen]TG[ArgNO27]FG[Sar][NMeL]YPC | 2000 |
| RF[Pen]TG[4Pal]FG[Sar][NMeL]YPC | 67 |
| RF[Pen]TG[3Pal]FG[Sar][NMeL]YPC | 640 |
| RF[Pen]TG[2Pal]FG[Sar][NMeL]YPC | 330 |
| RF[Pen]TG[Acc6393]FG[Sar][NMeL]YPC | 320 |
| RF[Pen]TG[Acc6018]FG[Sar][NMeL]YPC[NH2] | 110 |
| RF[Pen]TG[Tza]FG[Sar][NMeL]YPC | 840 |
| RFVTGHFG[Sar][NMeL]YPA | 1850 |

| | |
|---|---|
| QRFSTGHFGGLYPSNGP | 230000 |
| QRFKTGHFGGLYPENGP | 1200 |
| QRF[Dab]TGHFGGLYPENGP | 4800 |
| QRFKTGHFGGLYPDNGP | 10000 |
| QRF[Dab]TGHFGGLYPDNGP | 8400 |
| QRFETGHFGGLYPKNGP | 27000 |
| QRFETGHFGGLYP[Orn]NGP | 15000 |
| QRFETGHFGGLYP[Dab]NGP | 22000 |
| QRFETGHFGGLYP[db2Dap]NGP | 9600 |
| QRFDTGHFGGLYPKNGP | 1200 |
| QRFDTGHFGGLYP[Orn]NGP | 2700 |
| QRFDTGHFGGLYP[Dab]NGP | 17000 |
| QRFDTGHFGGLYP[db2Dap]NGP | 4500 |
| RF[Pen]TGHFG[Sar][NMeL]YPC | 31 |

The datasets cover a wide range in bioactivity (spanning up to 5 orders of magnitude), are structurally diverse in terms of varying sequence length, with circular peptides in dataset 2, and varying sequence similarity. Similarity is calculated as the number of identical pairs of residues divided by the total number of residues for a sequence and a reference, the reference taken as the first sequence in the set. Peptides in dataset 0 are high-similarity with 0.64 average, and with close to constant length of 12 with a small number of N-terminal insertions. Peptides in dataset 1 are low-similarity with 0.21 average, and a constant length of 9 and consist almost entirely of natural amino-acids with a few C-terminally O-methylated (OMe) peptides. Dataset 2 contains a small number of circular peptides (5/151), has an average similarity of 0.49 and sequence length of 16 residues. Similarity and sequence length distributions are plotted in Figure S3 (except the sequence length distribution for dataset 1 which is constant and equals 9) and summarized in Table S2.

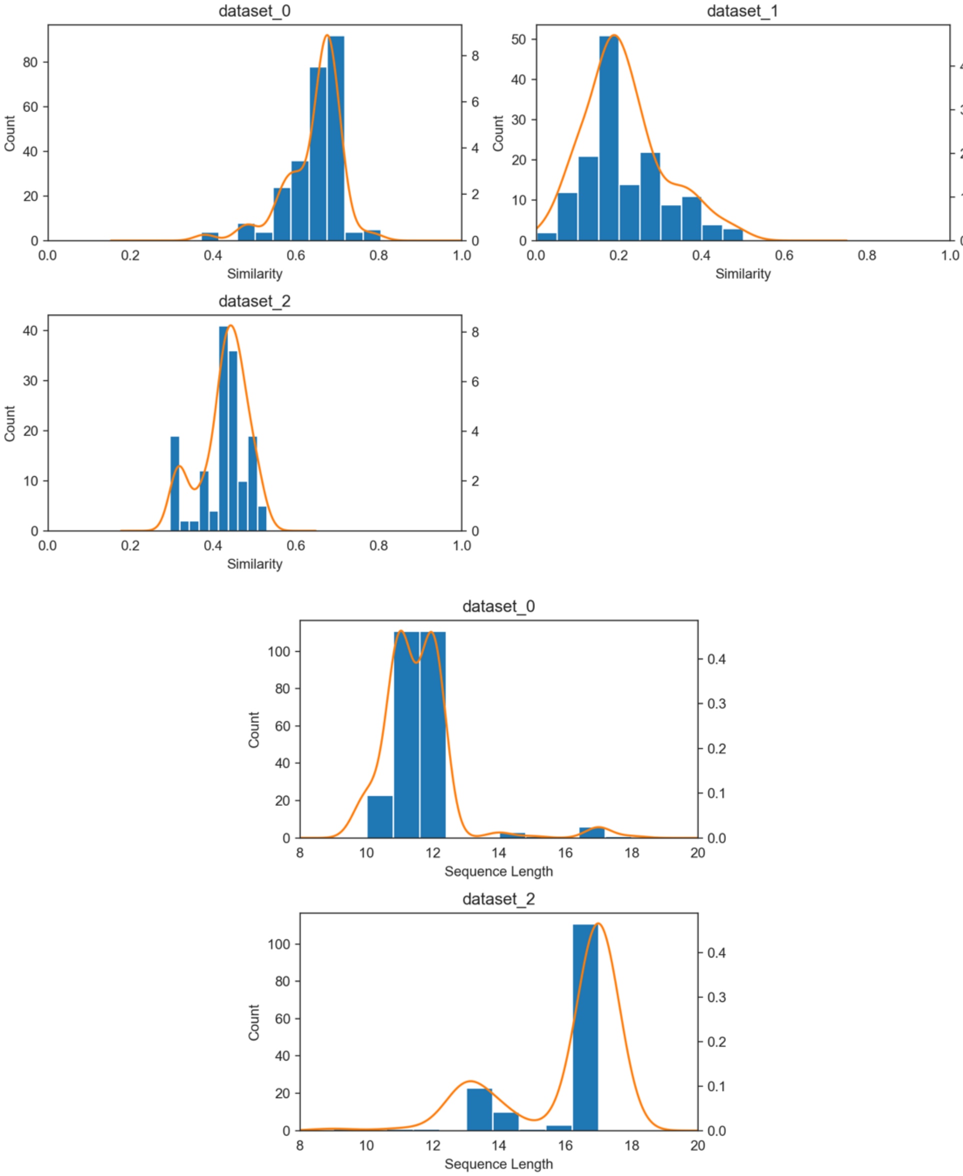


**Figure S3.** Similarity and sequence length distributions for the ChEMBL datasets 0, 1, 2 used in MSA benchmarking (see text for details).

To test the alignment of the ChEMBL datasets, we applied the ginsi and linsi alignment methods with default gap parameters, and with the larger values corresponding to the highest column scores for a dataset (Table 1). Also reported are the number of gaps in the MSA and the ROCS-based alignment score, calculated as the sum-of-pairs ROCS-score over all columns with the score for any residue aligned with a gap -10.

**Table S2.** Multiple Sequence Alignment for the ChEMBL datasets with g/linsi alignment methods, with default and the optimized gap parameters, defined as those with higher MSA-quality scores for simulated datasets. Similarity, sequence length and number of gaps are given as the average and range.

| ChEMBL-ID, Similarity, Sequence length | Method | gap_open | gap_extend | Number of gaps in MSA | Alignment Score |
|---|---|---|---|---|---|
| | linsi | 2.5 | 1.5 | 7 (1, 9) | 558 |
| CHEMBL1008221 | linsi | 1.53 | 0 | 7 (1, 9) | 583 |
| 0.64 (0.37 – 0.8) | ginsi | 2.5 | 0.5 | 7 (1, 9) | 583 |
| 12 (10 – 18) | ginsi | 1.53 | 0 | 8 (2, 10) | 560 |
| CHEMBL660105 | g/linsi | 3.0 | 3.0 | 0 (0, 0) | 390 |
| 0.21 (0 – 0.51) | | | | | |
| 9 (-) | g/linsi | 1.53 | 0 | 34, 43 | 84, -4 |
| | linsi | 2.5 | 1 | 4 (3, 11) | 451 |
| CHEMBL956457 | linsi | 1.53 | 0 | 4 (3, 11) | 451 |
| 0.43 (0.29 – 0.53) | ginsi | 2.5 | 0.5 | 4 (3, 11) | 451 |
| 16 (9 – 17) | ginsi | 1.53 | 0 | 4 (3, 11) | 451 |

The observation that the method ginsi is more sensitive to the choice of gap parameters, is evident with the CHEMBL1008221 dataset – which is not observed with the linsi method. Furthermore, the MSA using ginsi and the default gap parameters results in a lower alignment score, than with larger gap parameters (2.5, 0.5), which was also observed with simulated datasets. A striking difference in the alignment scores is observed for the low-similarity constant length CHEMBL660105 dataset – the MSAs with default gap parameters are extremely gappy, while using large gap parameters (3, 3) results in MSAs without gaps. The MSA alignment for the CHEMBL956457 dataset results in consistent alignment scores for all methods and gap parameters tested.

### Sequence alignment of simulated peptide dataset

To evaluate the performance of the ROCS-based alignment tool for the alignment of peptides, we used a simulation program ROSE[4] to generate the benchmark datasets. The idea was to generate peptides that resemble the original ChEMBL datasets in terms of sequence similarity and length distributions, for which we can evaluate the alignment quality given that their true-MSA is known. A set of 150 peptide sequences per dataset type were simulated by tweaking the parameters for substitution, insertion and deletion probabilities and lengths to achieve distributions comparable to those for the ChEMBL datasets (Figure S3). For the higher similarity datasets 0 and 2, sequences "TPQRARRRKKKG" and "QRFCTGHFGALYPCNP" were used as initial, while a random sequence with length 9 was used for the low-similarity dataset. Similarity and sequence lengths distributions of the 8 separate runs per dataset type are shown in Figure S4.

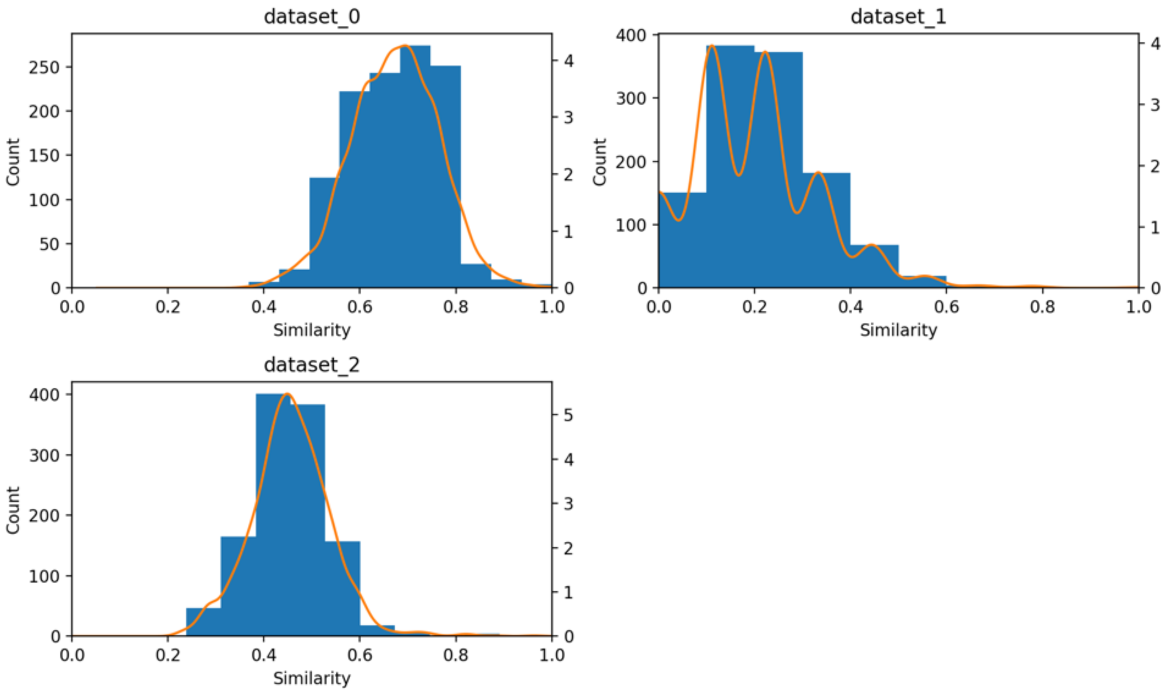

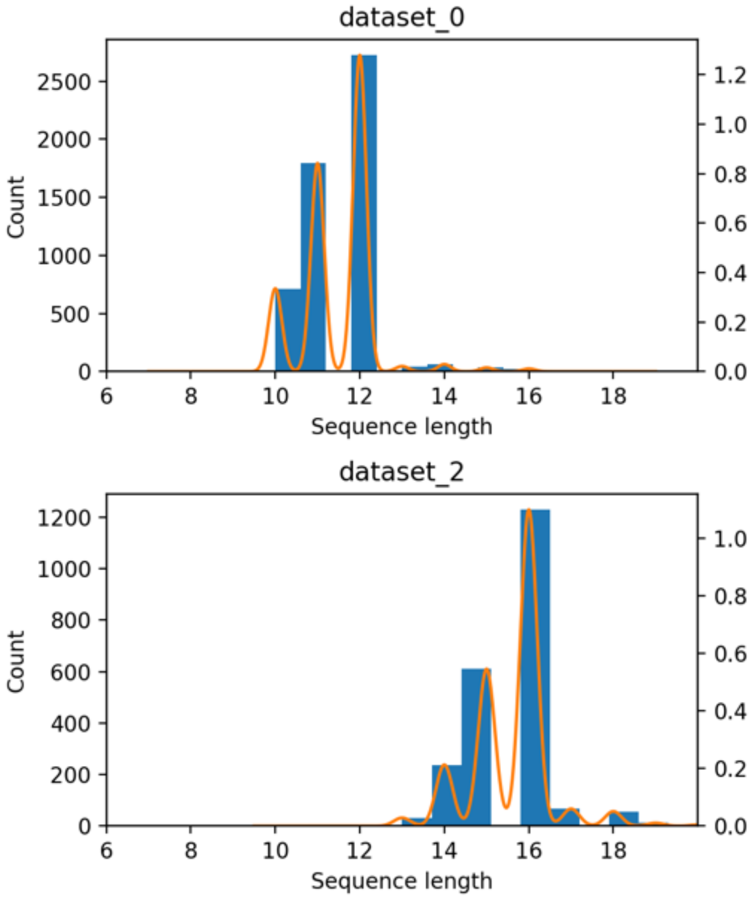


**Figure S4.** Similarity and Sequence length distributions for 8 runs of simulated datasets, aimed to resemble the ChEMBL datasets from Figure S3.

The comparison of the ChEMBL and the simulated datasets is further summarized in Table S2.

**Table S3.** Statistics for the simulated and ChEMBL datasets. Number of gaps for the ChEMBL dataset was estimated from an MSA generated with the method and gap parameters that resulted in the highest MSA-quality scores from simulations (see text).

| *Dataset* | *Sequence length* | *Similarity* | *Gaps in MSA** |
|---|---|---|---|
| 0 simulated | 11 (10, 16) | 0.67 (0.36 – 1.0) | 8 (2, 13) |
| 0 chembl | 12 (10, 18) | 0.64 (0.37 – 0.8) | 7 (1, 9) |
| 1 simulated | 9 – | 0.20 (0.0 – 0.78) | 0 – |
| 1 chembl | 9 – | 0.21 (0.0 – 0.51) | 0 – |
| 2 simulated | 16 (13, 20) | 0.46 (0.23 - 0.96) | 8 (3, 14) |
| 2 chembl | 16 (9, 17) | 0.43 (0.29 – 0.53) | 4 (3, 11) |

Common measures to evaluate the alignment quality are the Sum of Pairs (SPS) score and the Column Score (CS).[5] SPS score represents the fraction of pairs correctly reproduced in the tested MSA, compared to the reference MSA, calculated as

$$\mathrm{SPS} = \frac{\sum_{i=1}^{M} S_i}{\sum_{i=1}^{Mr} S_{ri}}\ ;\ S_i = \sum_{j=1,\, j\neq k}^{N} \sum_{j=1}^{N} p_{ijk}\ ;\ \ p_{ijk} = \begin{Bmatrix} 1\ , A_{ik} = A_{ik} \\ 0 \text{ otherwise} \end{Bmatrix}$$

For each pair of residues $A_{ij}$, $A_{ik}$, $p_{ijk}$ is 1 if residues are aligned in the reference alignment, 0 otherwise. The score for the *i-th* column of the alignment $S_i$, is calculated as the sum over all pairs of residues in a column, and the final SPS score as the sum over all columns normalized by the column sum in the reference MSA. Equivalently, the CS is calculated as a sum of identical columns between the test and the reference MSA, normalized by M, the number of columns in the tested MSA.

$$\mathrm{CS} = \sum_{i=1}^{M} \frac{C_i}{M} \; ; \; C_i = \begin{Bmatrix} 1 \text{ test} = \text{ref} \\ 0 \text{ otherwise} \end{Bmatrix}$$

The SPS score is a measure of the number of sequences correctly aligned, while the CS score measures the ability to align all the sequences correctly. The SPS and CS scores were calculated using the AlignStat R package for statistical comparison of MSAs.[6]

Different alignment strategies are implemented in the MAFFT program, including the faster progressive methods (FFT-NS-1/2) and the more accurate iterative methods which use either global or local (G/L/E-INS-I, referred to as ginsi, linsi, einsi further in text) algorithms in the pairwise alignment stage. Given that our datasets are highly variable in similarity and sequence length, we might expect different performance for global or local alignments. The gap opening and extension parameters were varied with values (1, 1.53, 2, 3, 5, 10) and (0, 0.5, 1, 2.5, 5), respectively, and the alignment methods tested were FFT-NS-2, ginsi, linsi, einsi. Interestingly, we observed a close to perfect SPS-scores (> 0.9) for all datasets with the g/linsi methods across the full range of gap parameters. For the *gappier* MSAs in datasets 0 and 2, linsi performed overall better than the global alignment (Figure S5), resulting in better scores over a wider range of gap parameters.

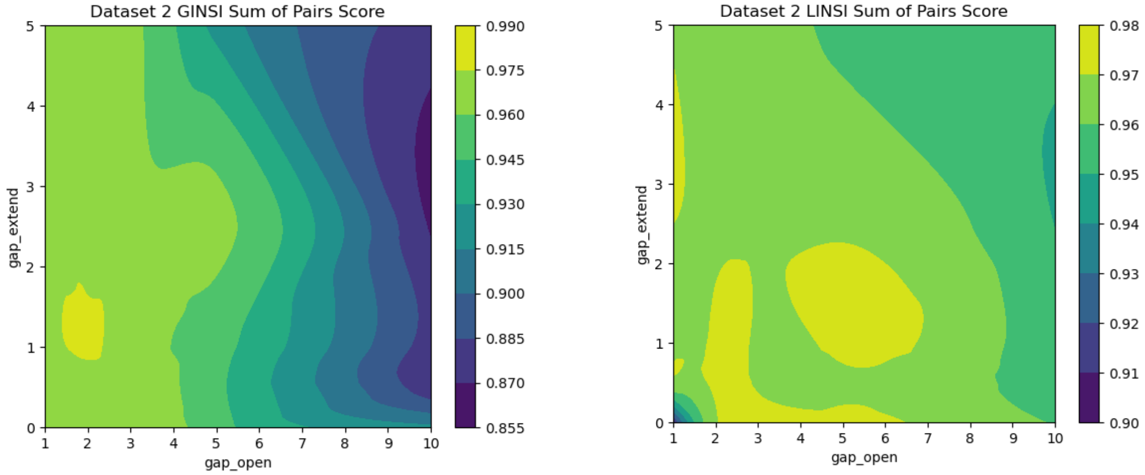


**Figure S5.** Comparison of the MSA quality SPS scores for dataset 2 (lower similarity, gappy dataset) for ginsi and linsi method.

On the other hand, the column scores were lower for the gappy-datasets 0, 2 and more sensitive to the gap parameters used (CS scores for the linsi method are plotted in Figure S6).

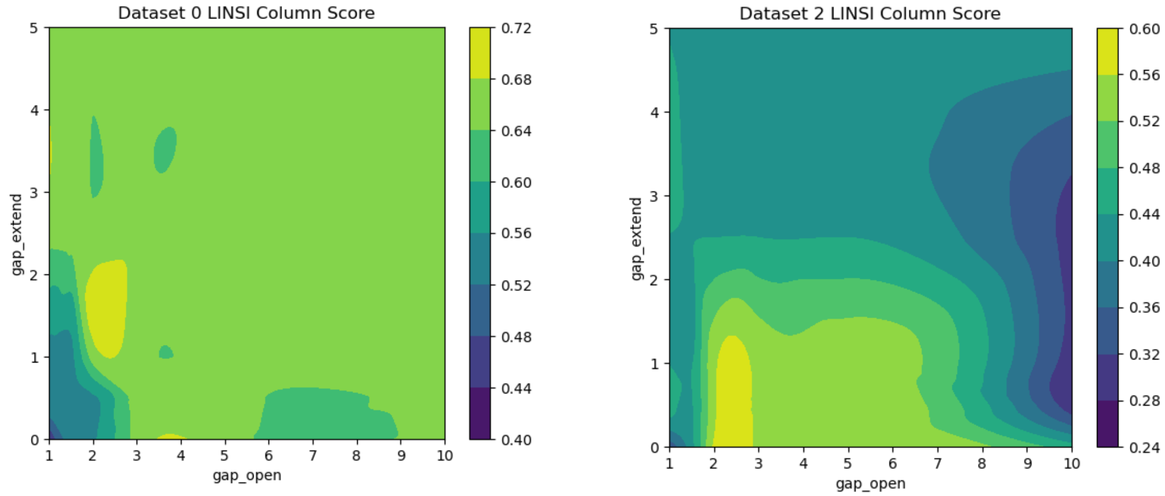


**Figure S6.** Column scores for datasets 0 and 2, where the test MSA was obtained with the linsi method.

The MSA using einsi-local alignment algorithm with a generalized affine gap, suitable for MSAs with long unalignable regions, also resulted in comparably high CS (0.6 - 0.7) and SPS scores (> 0.9), with larger values for gap parameters though (not shown here). Notably, for the low-similarity dataset 1 where simulated MSAs contained 0 gaps (see Table S2), the ginsi alignment was highly sensitive to the gap parameters used – for lower values, including the default for the MAFFT program (gap_open = 1.53, gap_extend = 0) the column scores were reduced to ~ 0.5, but otherwise achieved perfect alignment on the level of all sequences (CS = 1, Figure S7).

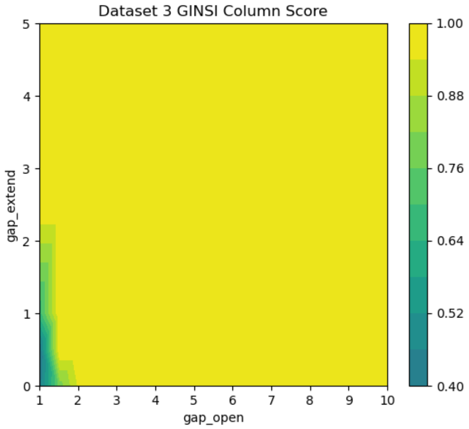


**Figure S7.** Column Score for the zero-gap dataset 1, using the ginsi method for alignment.

### PepSeA Visualization Tools

The visualization section consists of two parts, all written in Javascript: Visualization library and visualization API server. The browser-based visualization library is a module for transforming a HELM string to an SVG object. The library has two methods – one for transforming a HELM string to an object and another for transforming a parsed object to an SVG object. It supports custom color schema which can be imported before generating an SVG. The SVG-generation method takes additional parameters: Original Sequence – an array with chains of parsed HELM strings to compare monomers from data and highlight mismatches, Sequence Index – a sequence index to be included as metadata for rendered monomers (useful in case of many sequences), Linear – if true, generates an SVG with each chain as a line, or builds the chain as a circular structure with connections between monomers otherwise.

The Visualization API server is used for the representation of HELM string(s) and it is based on the Visualization library. It extends the Visualization library's parameters by the usePNG parameter, which makes the API return a representation in the PNG format.

https://doi.org/10.1186/s12859-016-1300-6.
